# Disassembly of Tau fibrils by the human Hsp70 disaggregation machinery generates small seeding-competent species

**DOI:** 10.1101/2019.12.16.876888

**Authors:** Eliana Nachman, Anne Wentink, Karine Madiona, Luc Bousset, Taxiarchis Katsinelos, Harm Kampinga, William A. McEwan, Thomas R. Jahn, Ronald Melki, Axel Mogk, Bernd Bukau, Carmen Nussbaum-Krammer

## Abstract

The accumulation of amyloid Tau aggregates is implicated in Alzheimer’s disease and other Tauopathies. Molecular chaperones are known for their function in maintaining protein homeostasis by preventing the formation or promoting the disaggregation of amorphous and amyloid protein aggregates. Here we show that an ATP-dependent human chaperone system disassembles Tau fibrils *in vitro*. This function is mediated by the core chaperone Hsc70, assisted by specific co-chaperones, in particular class B J-domain proteins and an Hsp110-type NEF. Recombinant fibrils assembled from all six Tau isoforms as well as Sarkosyl-resistant Tau aggregates extracted from cell culture were processed by the Hsp70 disaggregation machinery, demonstrating the ability of this machinery to recognize a broad range of Tau aggregates. Chaperone treatment released monomeric, and small oligomeric Tau species, which induced the aggregation of self-propagating Tau species in a Tau cell culture model. We infer from these results that the activity of the Hsp70 disaggregation machinery is a double-sided sword as it attempts to eliminate Tau amyloids but with the price of generating new seeds. The Hsp70 disaggregase therefore has a crucial function in the Tau propagation cycle, rendering it a potential drug target in Tauopathies.

## Introduction

Amyloid deposits are characteristic of various neurodegenerative diseases, such as Alzheimer’s (AD) and Parkinson’s disease (PD). Typically, symptoms surface only at advanced age indicating that a buffering system exists that prevents disease onset and amyloid formation earlier in life (Labbadia and Morimoto, 2015).

Disease-associated proteins aggregate into amyloid fibrils characterized by their highly ordered β-sheet structure (Chiti and Dobson, 2017). The monomeric form of these proteins populates conformations susceptible to aggregation, leading to the formation of a variety of assemblies of various molecular weights. Some have seeding propensities that trigger further aggregation into fibrils by templated incorporation of the monomeric form of the constituting protein in a conformation that is compatible with the fibril ends. This templated propagation of the amyloid structure is thought to be the basis for the prion-like spreading of pathological inclusions and toxicity in neurodegenerative diseases (Brundin et al., 2010).

Aggregation of the microtubule-associated protein tau (MAPT/Tau) is implicated in ∼20 different diseases termed Tauopathies, with AD being the most common form of dementia (Spillantini and Goedert, 2013). Tau is thus the most frequently aggregating protein in human neurodegenerative diseases. Under physiological conditions, Tau is highly soluble and consists of six alternatively spliced isoforms (Goedert et al., 1989). It binds and supports the assembly of microtubules that are vital for axonal transport in neurons (Mandelkow and Mandelkow, 2012). Under pathological conditions, the affinity of Tau to microtubules is reduced, either by disease-associated mutations or hyperphosphorylation (Ballatore et al., 2007; Biernat et al., 1993; Katsinelos et al., 2018). Detached Tau then forms aggregates in the cytoplasm that eventually evolve into fibrillar inclusions in affected neurons or glia cells (Spillantini and Goedert, 2013). In healthy cells, the homeostasis of Tau and other proteins is tightly controlled by a protein quality control network, including molecular chaperones (Klaips et al., 2018; Labbadia and Morimoto, 2015; Wentink et al., 2019). This quality control system protects the proteome by regulating the synthesis, folding, and trafficking of native proteins to their subcellular destination as well as the refolding and degradation of misfolded species. As such, molecular chaperones act at every step of the amyloid formation and clearance process (Balchin et al., 2016; Kampinga and Craig, 2010; Rosenzweig et al., 2019; Saibil, 2013; Wentink et al., 2019).

Numerous studies have linked molecular chaperone action to Tau aggregation both *in vitro* and *in vivo*. It was shown that Hsp70 family members, several J-domain protein co-chaperones, Hsp60, and the small Hsp Hsp27 delay Tau fibril formation (Abisambra et al., 2010; Mok et al., 2018; Patterson et al., 2011; Voss et al., 2012). Moreover, Hsp70 chaperones interact with oligomeric Tau and prevent further aggregation into fibrils (Patterson et al., 2011). So far, only inefficient disassembly of preformed Tau fibrils by Hsp70 activity has been observed (Patterson et al., 2011; Voss et al., 2012). Our work and other studies demonstrated that Hsp70 disaggregation activity strongly relies on co-chaperone action (Kampinga and Craig, 2010; Rosenzweig et al., 2019). J-domain proteins deliver clients to Hsp70 by pre-selecting them and activating their ATP hydrolysis-dependent binding into the substrate binding pocket of Hsp70, thereby determining substrate specificity of the machinery. Nucleotide exchange factors (NEFs) regulate the lifetime of the Hsp70-substrate complexes, which determines substrate fate, such as refolding, transfer to other chaperone systems or handover to the degradation machinery (Bracher and Verghese, 2015). To date, the correct composition of chaperones and co-chaperone combinations that efficiently dissolves Tau fibrils is unknown.

Here we demonstrate that the human Hsp70 disaggregation machinery, referred to hereafter as “Hsp70 disaggregase”, has the capacity to disassemble amyloid Tau fibrils *in vitro*. The Hsp70 disaggregase is an ATP-dependent chaperone system, which is comprised of the constitutively expressed Hsp70 family member Hsc70, the J-domain proteins DnaJB1 or DnaJB4, and Apg2, an Hsp110-type NEF. Both, recombinant fibrils of all six Tau isoforms and Sarkosyl-resistant Tau aggregates extracted from a cell culture model, could be processed by this chaperone system. We further show that class B J-domain proteins are essential for this activity and that there is partial redundancy within this class of chaperones, while class A J-domain proteins were not able to support Hsp70 disaggregase function. The disaggregation reaction produced monomeric Tau as well as small oligomers. Tau species liberated by the disaggregation reaction were seeding-competent and induced self-propagating Tau species in a biosensor cell line for Tau aggregation.

This study shows that Hsp70 disaggregase activity can be extended to the most prevalent neurodegenerative diseases involving Tau. Although almost completely depolymerized to monomers, the fraction of Tau, which was released from amyloid fibrils by chaperone action, was still seeding-competent. As the generation of seeding-competent species might boost the prion-like propagation of amyloid Tau aggregates, it needs to be examined whether chaperone-mediated Tau disaggregation may exacerbate the associated neurotoxicity *in vivo*.

## Results

### The human Hsp70 disaggregase disassembles recombinant Tau fibrils *in vitro*

To investigate whether Tau fibrils can be disassembled by the human Hsp70 disaggregase, we performed *in vitro* disaggregation assays (Gao et al., 2015) (Fig. 1A). Recombinant 1N3R Tau, harboring one N-terminal insertion and three microtubule binding repeats (Fig. S1), was assembled into fibrils and fibril formation was verified by negative stain transmission electron microscopy (TEM) (Fig. 1B). Tau fibrils were treated with the human Hsp70 disaggregation machinery (Hsc70, DnaJB1, Apg2) and subsequently centrifuged in order to separate larger fibrils from liberated smaller oligomers and monomers. The amount of Tau in supernatant (S) and pellet (P) fractions was analyzed by SDS-PAGE and immunoblotting (Fig. 1C). In the presence of the three chaperones and ATP more than 40% of 1N3R Tau was detected in the supernatant fraction (Fig. 1C and 1D). In contrast, in the absence of ATP, the chaperone mix did not confer any significant disaggregation activity (Fig. 1C and 1D). The three components of the Hsp70 disaggregation machinery were also added individually and in all possible combinations to test their respective contribution to fibril disassembly (Fig. 1E and 1F). Treatment with single or pairwise combinations of chaperones did not promote a shift of Tau to the supernatant fraction, except Hsc70 together with DnaJB1, which resulted in relocation of ∼28% of Tau to the supernatant (Fig. 1E and 1F). Yet, the combination of all three chaperones was required for a most efficient disaggregation reaction leading to ∼43% disassembly (Fig. 1E and 1F), indicating a critical role of the HSP110-type NEF in Tau fibril disaggregation. In conclusion, the human Hsc70/DnaJB1/Apg2 disaggregation machinery efficiently disassembles a significant fraction of Tau fibrils *in vitro*.

**Figure 1.**
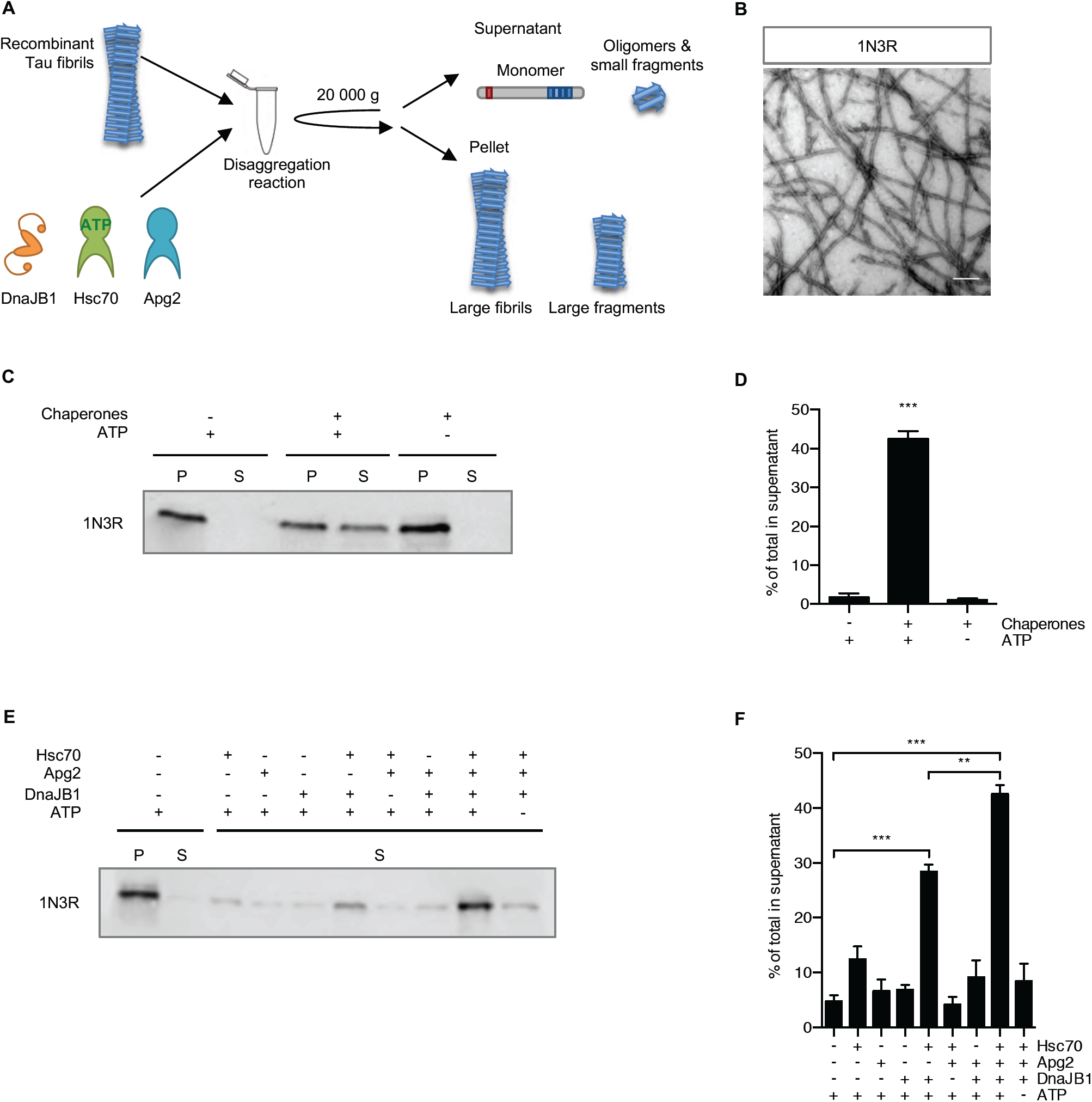
The human Hsp70 disaggregation machinery disassembles recombinant Tau fibrils. **(A)** Experimental setup of the disaggregation reaction and subsequent sedimentation assay. Tau fibrils were treated with chaperones for 20 h at 30 °C. Large fibrils are separated from smaller species and monomers by centrifugation at 20 000 g. **(B)** Electron micrograph of negatively stained fibrils from recombinant 1N3R Tau assembled *in vitro*. Scale bar = 250 nm. **(C)** Fibrils (2 *µ*M) were incubated with chaperones (4 *µ*M Hsc70, 2 *µ*M DnaJB1, 0.2 *µ*M Apg2) ± ATP for 20 h at 30 °C. Supernatant (S) and pellet (P) fractions were separated by centrifugation (20 000 g) and analyzed by immunoblotting. **(D)** Densitometric quantification of Tau in S fractions compared to total (S+P) of each sample of the western blot shown in (C). n = 3, mean ± SEM. Statistical analysis was done using a one-way ANOVA with Bonferroni’s multiple comparison test. *** p ≤ 0.001. **(E)** Tau fibrils were treated with either individual chaperones of the disaggregation machinery or all possible combinations, respectively. S and P fractions were separated by centrifugation and Tau levels were analyzed by immunoblotting. Supernatant fractions of a representative experiment are shown. **(F)** Densitometric quantification of Tau in supernatant fractions compared to total (S+P) of each sample of the experiment shown in (E). n = 3, mean ± SEM. Statistical analysis was done using a one-way ANOVA with Bonferroni’s multiple comparison test. For clarity only the significances to the -Chap +ATP condition and between Hsc70, DnaB1 + ATP in the presence or absence of Apg2 are indicated. ** p ≤ 0.01, *** p ≤ 0.001.

### All six Tau isoforms can be disassembled by the disaggregation machinery

Human Tau has six different isoforms that are generated by alternative splicing (Goedert et al., 1989) (Fig. S1). Whereas all isoforms were found in aggregates isolated from AD patients’ brains, there are also isoform-specific Tauopathies where amyloid deposits consist exclusively of either 3R or 4R Tau isoforms (Spillantini and Goedert, 2013). For example, Tau filaments in Pick’s disease contain only 3R Tau, while progressive supranuclear palsy (PSP) is characterized by fibrils made entirely of 4R isoforms.

To test whether all six isoforms are substrates for the disaggregation machinery, recombinant fibrils of the other five Tau isoforms (0N3R, 2N3R, 0N4R, 1N4R, and 2N4R) (Fig. 2A) were subjected to disaggregation and the reaction products were analyzed by differential centrifugation (Fig. 2B). Similar to 1N3R fibrils, fibrils formed by all other Tau isoforms could be disassembled by the human disaggregation machinery, although with varying efficiencies (Fig. 2B and 2C). In general, all 3R isoforms displayed higher disaggregation rates compared to their 4R counterparts, with 0N3R Tau fibrils being most efficiently disassembled (53%) and 0N4R fibrils being most resistant to chaperone mediated disaggregation (11%). Overall these results show that the Hsp70 disaggregase exhibits disaggregation activity towards all Tau variants and is not limited to fibrils assembled from a certain Tau isoform.

**Figure 2.**
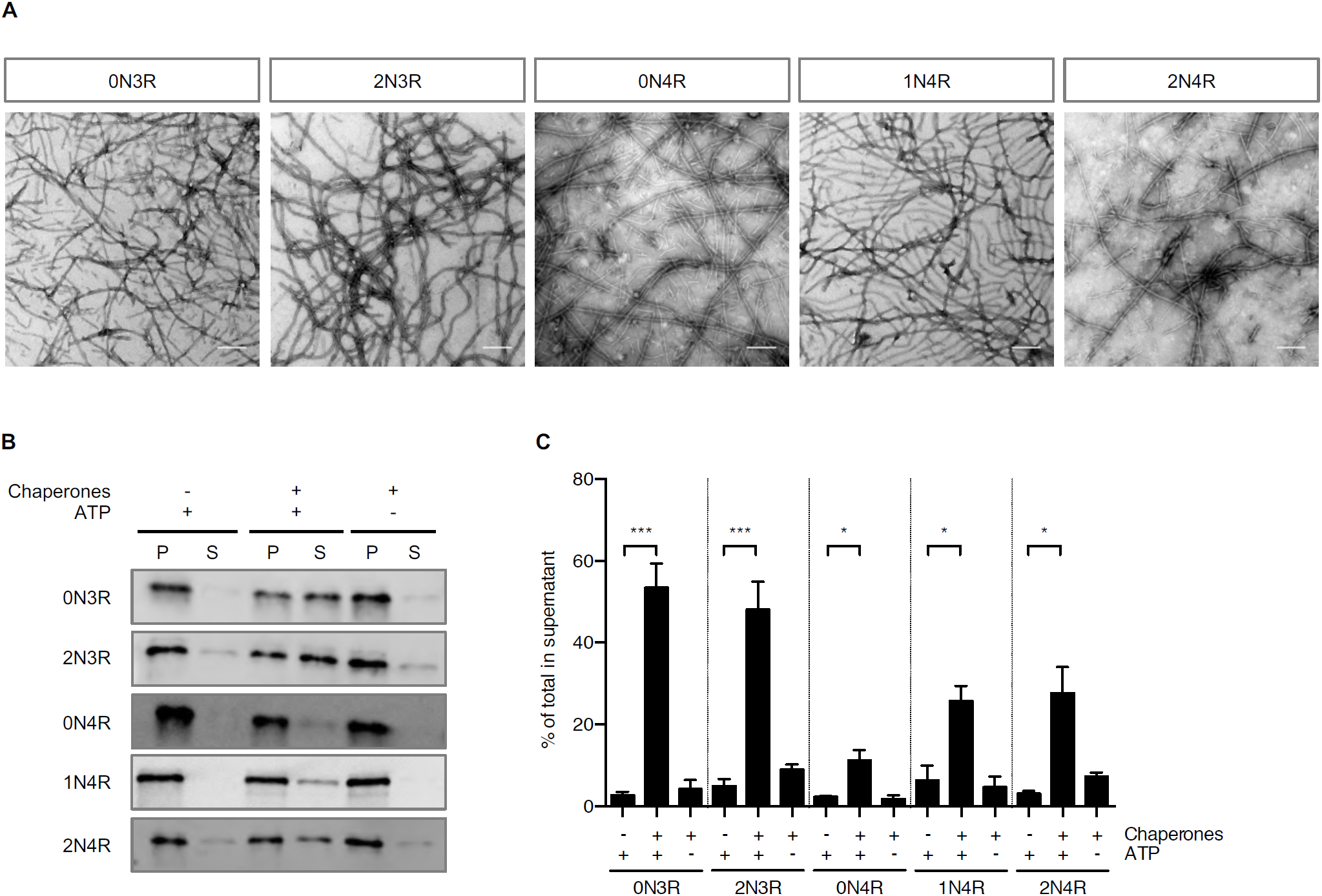
All six Tau isoforms are substrates for the disaggregation machinery. **(A)** Electron micrographs of negatively stained fibrils from recombinant Tau isoforms aggregated *in vitro*. Scale bar = 250 nm. **(B)** Recombinant fibrils formed from all five Tau isoforms were incubated with the Hsc70, DnaJB1, Apg2 disaggregation machinery for 20 h at 30 °C. S and P fractions were separated by centrifugation and analyzed by Western blotting. **(C)** Densitometric quantification of Tau levels in the experiment shown in (B). n = 3, mean ± SEM. One-way ANOVA with Bonferroni’s multiple comparison test. Significances are shown compared to -Chap +ATP condition for each isoform, respectively. * p ≤ 0.05, *** p ≤ 0.001.

### Detergent-insoluble Tau extracted from cell culture can be disaggregated

Amyloid aggregates formed *in vivo* might possess different properties than *in vitro* aggregated fibrils as post-translational modifications or co-aggregation with other endogenous proteins could affect the overall structural arrangement (Fichou et al., 2019). In order to investigate whether Tau aggregates formed in cells are clients of the human Hsp70 disaggregation machinery we made use of a HEK293 cell model of Tau aggregation (McEwan et al., 2017). This cell line constitutively overexpresses Venus-tagged full-length P301S mutant 0N4R Tau (0N4R TauP301S-Venus), which remains Sarkosyl-soluble under normal growth conditions and was successfully used as biosensor for Tau seeding (McEwan et al., 2017). After treating the cells with recombinant fibrils from the 1N4R Tau isoform, we specifically enriched the seeded cells through flow cytometry sorting and expanded them to provide a source of Tau aggregates that were formed in cells (Fig. 3A). The Sarkosyl-resistant material was extracted from the seeded cells and subjected to *in vitro* disaggregation assays. In the presence of Hsc70, DnaJB1, Apg2, and ATP 30% of the TauP301S-Venus was recovered in the supernatant fraction following centrifugation (Fig. 3B and 3C). This result demonstrates that the human Hsp70 disaggregation machinery is capable of disassembling Sarkosyl-insoluble Tau extracted from a human cell line.

**Figure 3.**
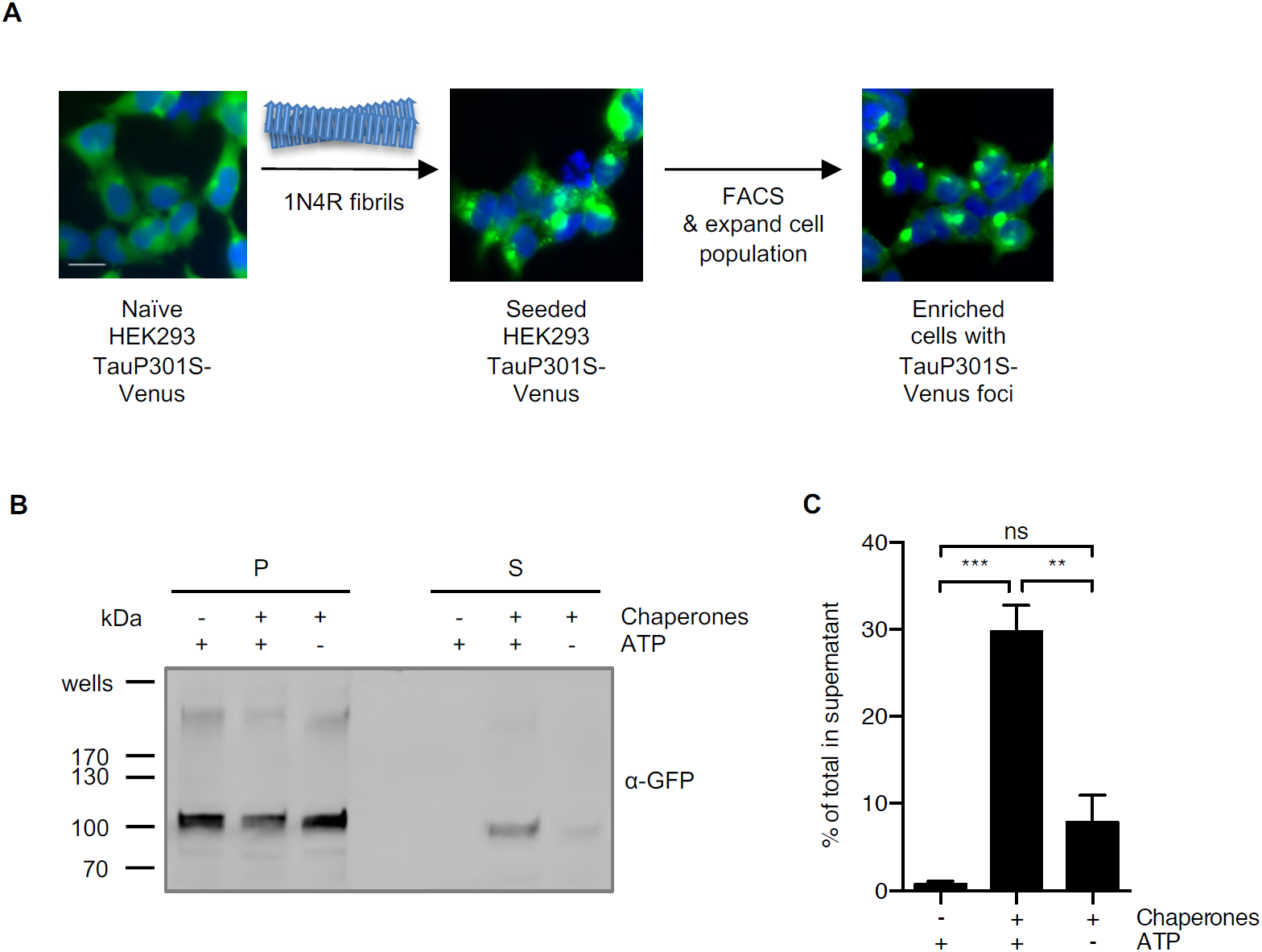
Sarkosyl-insoluble TauP301S-Venus aggregated in a cell culture model is a substrate for the disaggregation machinery. **(A)** A HEK293 0N4R TauP301S-Venus cell line was used to generate a cell population with Tau aggregates as outlined in the depicted workflow. Naïve cells were seeded with 1N4R fibrils followed by the enrichment of foci containing cells by FACS. A representative image of the seeded cell population after the FACS enrichment is shown. **(B)** Sarkosyl-insoluble TauP301S-Venus material extracted from the seeded HEK293 cell population was subjected to *in vitro* disaggregation assays with the recombinant human Hsp70 disaggregation machinery (Hsc70, DnaJB1, Apg2) ± ATP for 20 h at 30 °C. Supernatant (S) and pellet (P) fractions were separated by centrifugation at 337 000 g and analyzed by immunoblotting with an α-GFP antibody detecting TauP301S-Venus. **(C)** Densitometric quantification of TauP301S-Venus in S fractions compared to total (S+P) of the western blot shown in (B). n = 3, mean ± SEM. One-way ANOVA with Bonferroni’s multiple comparison test. ns = not significant, *** p ≤ 0.001.

### Class B J-domain proteins mediate disaggregation

Hsp70 substrate specificity is mediated by J-domain proteins that recognize chaperone clients and deliver them to Hsp70 (Kampinga and Craig, 2010). Humans encode more than 40 different J-domain proteins, subdivided into structural classes A, B, and C, with distinct substrate specificities and cellular localization (Kampinga and Craig, 2010; Rosenzweig et al., 2019). Several class A and B J-domain proteins are differentially regulated in the brain both during aging and in neurodegenerative diseases (Brehme et al., 2014). In particular, the class A co-chaperone DnaJA1 as well as the class B co-chaperone DnaJB4 are upregulated in patients with AD, PD, and Huntington’s disease (HD) compared to age-matched controls (Brehme et al., 2014). These findings point to a potential role of these co-chaperones in regulating proteostasis in the context of protein misfolding diseases.

Therefore, we investigated whether these J-domain proteins could also serve in a complex with Hsc70 and Apg2 to disaggregate Tau fibrils (Fig. 4A). The class B J-domain protein DnaJB4 was equally capable of promoting disaggregation of recombinant 1N3R Tau fibrils as DnaJB1, both shifting ∼50% of Tau to the supernatant fraction (Fig. 4A and 4B). Although 1N4R fibrils were again less susceptible to disassembly by the Hsp70 disaggregation machinery than 1N3R fibrils, DnaJB4 could also substitute for DnaJB1 and mediated ∼36% disaggregation compared to ∼26% for DnaJB1 (Fig. 4A and 4C). The two class A J-domain proteins tested here, DnaJA1 and DnaJA2, did not enable disaggregation of either 1N3R or 1N4R Tau fibrils (Fig. 4A-4C). Moreover, DnaJA2 did not promote disaggregation of any other Tau isoform (Fig. S2A). In conclusion, the class B J-domain protein DnaJB4, which is closely related to DnaJB1 (Fig. S2B and S2C), also enabled efficient disaggregation of Tau amyloid fibrils while the class A J-domain proteins could not assist their disassembly.

**Figure 4.**
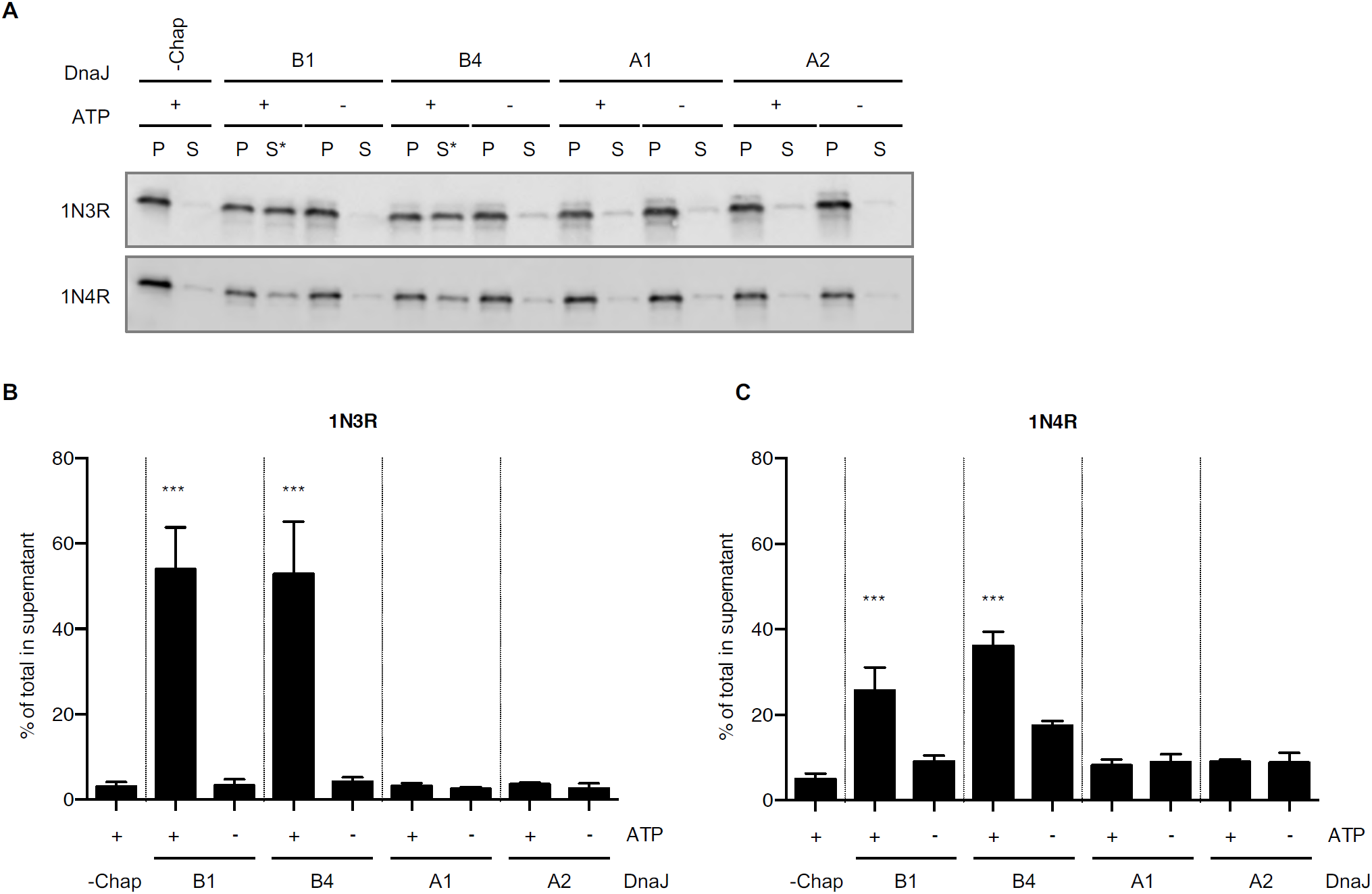
Class B J-domain proteins mediate Tau disaggregation. **(A)** Tau fibrils made up from 1N3R or 1N4R were treated with Hsc70, Apg2, and different Hsp40 class A and class B J-domain proteins, respectively. Subsequently, S and P fractions were separated by centrifugation and Tau levels were analyzed by Western blotting. Asterisk (*) indicate successful disaggregation reactions. **(B)** Densitometric quantification of 1N3R Tau levels in S fractions compared to total (S+P) as shown in (A). n = 3, mean ± SEM. One-way ANOVA with Bonferroni’s multiple comparison test, only significances to -Chap +ATP condition are indicated. *** p ≤ 0.001. **(C)** Densitometric quantification of 1N4R Tau levels in S fractions compared to total (S+P) as shown in (A). n = 3, mean ± SEM. One-way ANOVA with Bonferroni’s multiple comparison test, only significances to -Chap +ATP condition are indicated. *** p ≤ 0.001.

### Tau disaggregation yields monomeric and small oligomeric seeding-competent species

The products of the disaggregation reaction could consist of multiple protein species, such as monomers, small oligomers and other fibril fragments with intermediate lengths. Furthermore, it is not yet clear whether chaperone-mediated disaggregation of amyloid fibrils is advantageous or disadvantageous. A complete resolubilization and refolding of Tau into monomers is considered beneficial, while the production and accumulation of fibrillar intermediates is considered disadvantageous, as the latter could contribute to the propagation of Tau aggregates through the continuous production of new seeds. Therefore, it is important to analyze the products of the disaggregation reaction more closely.

In order to monitor the disaggregation dynamics, we determined the quantity of amyloid structures during the course of chaperone-mediated disaggregation, by measuring the fluorescence of the amyloid-specific dye ThT over time. Incubation of both 1N3R (Fig. 5A) and 1N4R (Fig. 5B) Tau fibrils with the disaggregation machinery and ATP resulted in a decrease in ThT fluorescence of 25% and 10% within 4 h, respectively. After that, the ThT fluorescence plateaued over the course of the measurement. In the absence of ATP, the ThT fluorescence remained stable over time. This observation reflected the results obtained by differential centrifugation for the two Tau isoforms (Fig. 1C and 1D, Fig. 2B and 2C). However, the disaggregation efficiency determined by reduction in ThT fluorescence was in general lower compared to the efficiencies obtained by the sedimentation assay and subsequent immunoblotting. This is likely due to the generation of ThT positive oligomers or small fibrillar fragments generated during the disaggregation reaction, that are small enough to remain in the supernatant fraction after centrifugation.

**Figure 5.**
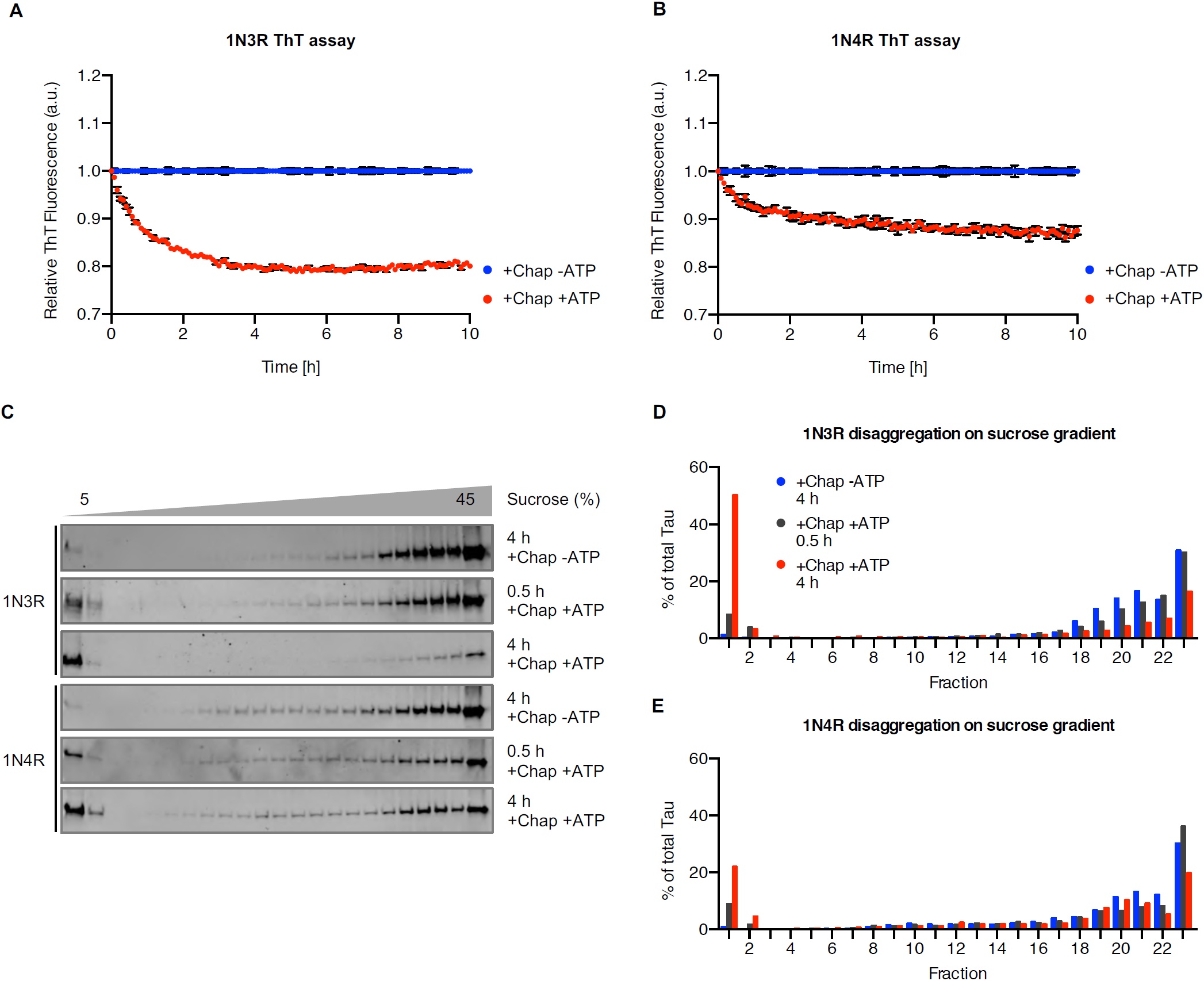
The disaggregation reaction generates low molecular weight species. **(A)** 1N3R and **(B)** 1N4R Tau fibrils were treated with chaperones in the presence or absence of ATP. Thioflavin (ThT) fluorescence was monitored over time as a readout of amyloid content. n = 3, mean ± SEM. **(C)** Tau fibrils were treated with the disaggregation machinery for 30 min or 4 h at 30 °C and subsequently centrifuged over a 5-45 % sucrose gradient. Fractions were collected manually, and Tau levels were analyzed by immunoblotting. **(D)** Densitometric quantification of 1N3R disaggregation and 1N4R disaggregation **(E)** of the experiment shown in (C). The amount in each fraction was calculated as a percentage of the total amount of Tau in all fractions.

To characterize the composition of the disaggregated Tau material in more detail, we performed rate-zonal centrifugation, in which Tau assemblies are separated according to their size. We first performed control experiments with untreated monomeric and fibrillar Tau, respectively (Fig. S3). After 3 h of centrifugation over a 5-45% sucrose gradient, monomeric recombinant 1N3R and 1N4R Tau remained in the first two low-density fractions, while untreated fibrils migrated to the higher density fractions (Fig. S3). Next, 1N3R and 1N4R Tau fibrils were analyzed after 30 min and 4 h of chaperone treatment. These time points were chosen based on the disaggregation kinetics followed using ThT binding, reflecting the intermediate and end points of the disaggregation reaction, respectively (Fig. 5A and 5B).

In the presence of the Hsc70, DnaJB1, Apg2, and ATP, an increasing amount of Tau was detected in the low-density fractions over the course of the disaggregation reaction, while the amount of Tau in all middle and high-density fractions was reduced (Fig. 5C-5E). Fibrils incubated for 4 h with chaperones but without ATP moved to the higher density fractions and no shift of Tau to lower density fractions was observed (Fig. 5C-5E). We did not detect any buildup of Tau in intermediate fractions during the disaggregation reaction (Fig. 5D and 5E). This finding was also in line with our EM analysis where we did not observe widespread fibril fragmentation (Fig. S4). Instead, most of the Tau protein liberated by the disaggregation machinery shifted to the low-density fractions in the sucrose gradients, suggesting that predominantly monomeric or small oligomeric Tau species were released.

To further characterize these low molecular weight Tau species, we next subjected Tau fibrils with and without chaperone treatment to sequential centrifugation (Fig. 6A). In untreated 1N4R Tau fibril preparations about 6% of Tau was found in the 20 000 g supernatant (Fig. 2B). However, these Tau species sedimented at 337 000 g while the Tau material that was released by disaggregation remained in the supernatant hinting towards a smaller particle size of these species (Fig. 6B). Hence, the 337 000 g supernatant contains Tau species specifically produced by the action of the Hsp70 disaggregation machinery, which are not present without chaperone treatment. Tris-acetate-SDS-PAGE revealed that the disaggregated material contained monomeric as well as oligomeric Tau species with an apparent molecular weight compatible with that of dimeric and tetrameric Tau (Fig. 6C). The latter migrated as distinct bands with apparent molecular weights of 100 kDa and 200 kDa in the gel and could not be dissolved by incubating in 2% SDS at 22 °C or 95 °C (Fig. 6C).

**Figure 6.**
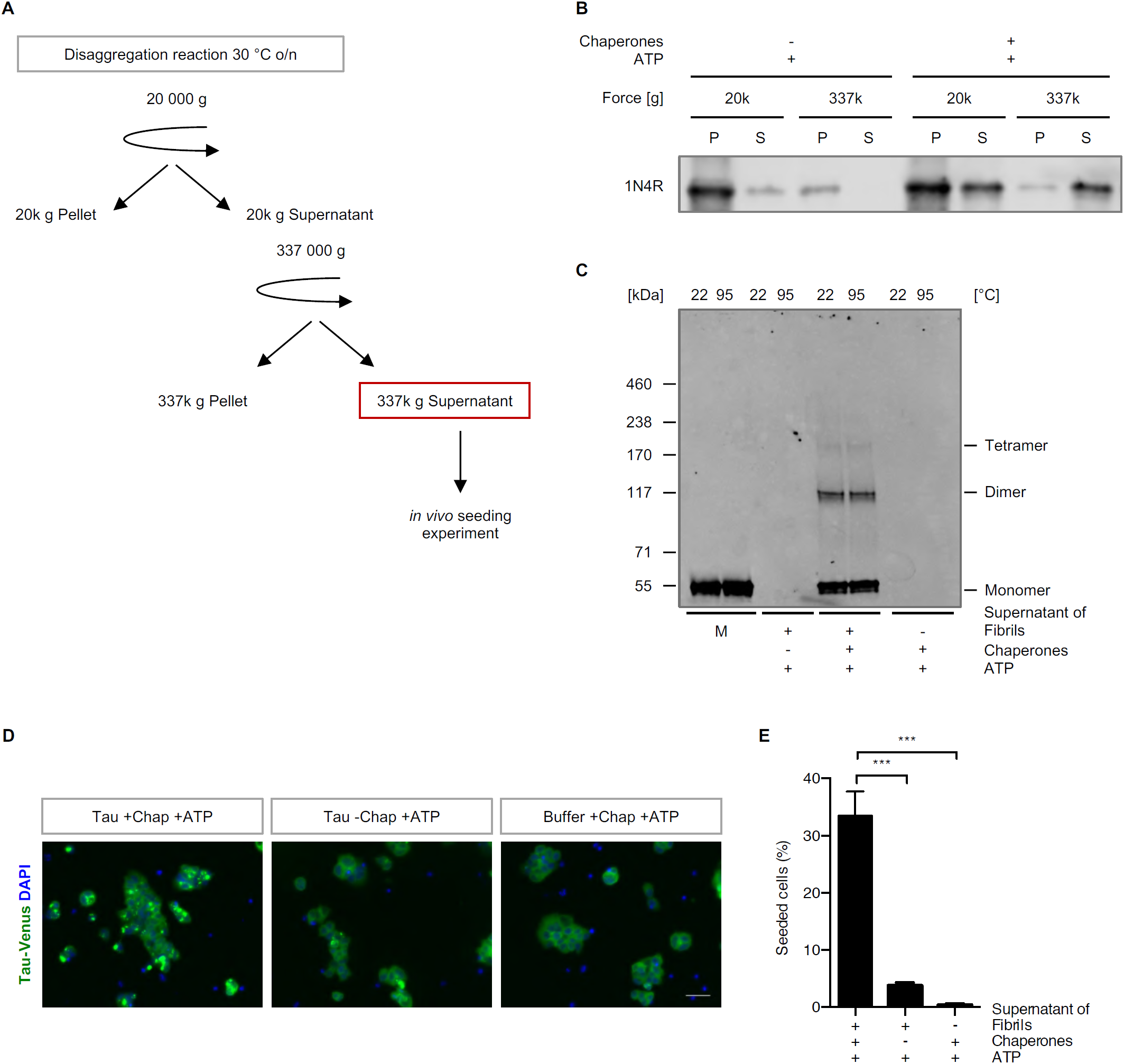
The disaggregation reaction liberates seeding-competent Tau species. **(A)** Experimental set-up of the *in vivo* seeding assay. Differential centrifugation of 20 000 g followed by ultracentrifugation at 337 000 g was applied in order to isolate the Tau material which was liberated by the action of the Hsp70 disaggregation machinery. The 337 000 g S fractions were tested for their seeding capacity in a HEK293 cell culture model for Tau aggregation. **(B)** Tau levels in the S and P fractions that were collected during the differential centrifugation steps shown in (A) were analyzed by immunoblotting. **(C)** The 337 000 g S fractions of disaggregated Tau and monomeric Tau were incubated with 2% SDS at room temperature or 95 °C and run on a Tris-acetate-SDS-PAGE. The samples were analyzed by immunoblotting. **(D)** Maximum intensity projections of fluorescence microscopy z-stacks of TauP301S-Venus HEK293 cells seeded with the 337 000 g supernatant fraction after the disaggregation reaction. Scale bar = 50 *µ*m. **(E)** Quantification of cells containing TauP301S-Venus foci. n = 3 replicates with 288 - 460 cells per condition in each replicate, mean ± SEM. Statistical analysis was performed using a one-way ANOVA with Bonferroni’s multiple comparison test. *** p ≤ 0.001.

Finally, we evaluated the seeding propensity of this fraction of Tau released by the Hsp70 disaggregase in the TauP301S-Venus HEK293 cell line (McEwan et al., 2017). The TauP301S-Venus expressing HEK293 cells were exposed to this material for 24 h and the percentage of foci-containing cells was evaluated by fluorescence microscopy. The 337 000 g supernatant of Tau fibrils incubated with the disaggregation machinery and ATP triggered foci formation in ∼33% of cells. None of the following, the supernatant of untreated fibrils, the chaperones of the disaggregation machinery and ATP, buffer with ATP (Fig. 6D and 6E), and naïve monomeric Tau (Fig. S5A) induced foci formation. We further assessed the longitudinal propagation of the seeded Tau species, i.e. the inheritance of Tau foci in cells transiently exposed to the Tau released by chaperone action. These Tau foci persisted in daughter cells during subsequent cell passages (Fig. S5B), demonstrating that Tau species generated by the Hsp70 disaggregation machinery were not only seeding-competent but were also able to induce self-propagating aggregate species.

In conclusion, Tau fibril disassembly by the human Hsp70 disaggregation machinery did liberate monomeric, as well as dimeric and tetrameric Tau species, which were seeding-competent and induced self-propagating Tau conformers in a HEK293 cell culture model for Tau aggregation.

## Discussion

It is well established that the cellular network of molecular chaperones assists in all aspects of protein quality control, from folding of newly synthesized peptides to the disassembly of protein aggregates and degradation of terminally misfolded proteins (Klaips et al., 2018; Labbadia and Morimoto, 2015; Wentink et al., 2019). Chaperones thereby affect many disease states, which is why chaperone-based therapies could be a promising treatment approach. However, it remains poorly understood to what extent chaperones are capable of disassembling already existing amyloids, given their high thermodynamic stability (Baldwin et al., 2011). Only for α-synuclein and HTTExon1Q_48_ it has been established that the human Hsp70 machinery is able to disassemble preformed fibrils *in vitro* (Gao et al., 2015; Scior et al., 2018). Here we investigate the broader role of this machinery in amyloid biology by testing its potential to process aggregates of amyloidogenic Tau isolated from cells or produced *in vitro* and by characterizing more precisely the products of chaperone-mediated Tau disaggregation. This is particularly important because Tau aggregation is central to the most prevalent human neurodegenerative diseases, including AD, and also plays a role in traumatic brain injuries (McKee et al., 2013; Spillantini and Goedert, 2013).

We show that the Hsp70 disaggregation machinery is capable of disassembling *in vitro* aggregated Tau amyloid fibrils as well as Sarkosyl-resistant Tau aggregates formed in cells. Tau disaggregation resulted in a rapid accumulation of low molecular weight Tau species (Fig. 5C and 6C). Further characterization of the liberated Tau pool revealed that it contained mostly monomeric and also some oligomeric species with apparent molecular weights of ∼100 kDa and ∼200 kDa, compatible with that of dimeric and tetrameric Tau, respectively. Intriguingly, this material was still seeding-competent as it induced longitudinal self-propagating aggregates of a stably expressed full-length Tau reporter in a HEK293 cell culture model, implying that chaperone-mediated Tau disaggregation is not per se beneficial, but may be involved in the prion-like propagation of Tau pathology.

Amyloid fibrils share a common core structure consisting of a characteristic β-sheet rich conformation (Jahn et al., 2010). Although exhibiting a very similar architecture, fibrils composed of α-synuclein, HTTExon1Q_48_, and Tau will display different surface properties, as they do not share any sequence homology (Knowles et al., 2014; Melki, 2018; Tycko, 2015). Nevertheless, the Hsp70 disaggregation machinery is able to disassemble amyloid fibrils composed of each of these proteins albeit with varying efficiencies, highlighting the versatility of this chaperone system to process various amyloid substrates.

Still, despite overall structural similarity, conformational variations of the amyloid structure formed by a given protein including Tau are known to exist and to affect the pathology of the associated disease (Knowles et al., 2014; Melki, 2018; Tycko, 2015). This variability could make some fibrils more resistant to chaperone action. We indeed observed differences in disaggregation efficiencies between the six distinct Tau isoforms. Fibrils comprised of 0N4R Tau were most resistant to disaggregation resulting in only 10% disaggregated material compared to up to 64% obtained with the other isoforms. The 0N4R Tau isoform remained also intact after the addition of the Hsp70 disaggregation complex in another study (Mok et al., 2018). Fibrils assembled from 3R Tau isoforms were disassembled to a greater extent than those made of the 4R isoforms (Fig. 1D and 2C). *In vitro* assembled fibrils from 2N3R and 2N4R Tau vary in their architecture (Zhang et al., 2019). The ordered core of 2N3R Tau fibrils comprises the R3 repeat of two parallel Tau molecules, whereas 2N4R Tau fibrils adopt several conformations with a core consisting of R2 and R3 β-strands of the same molecule (Zhang et al., 2019). Intriguingly, these differences in fibril architecture may lead to a different stability of 3R and 4R Tau fibrils, which may explain their varying susceptibility to the human Hsp70 disaggregation machinery. Alternatively, or additionally, the kinetics of chaperone binding may vary between isoforms, leading to different disaggregation efficiencies.

We never observed a disaggregation efficiency greater than 64%. This could be due to a mixture of different Tau conformers in our fibril preparation that is subjected to disaggregation. A subset thereof might be readily disaggregated while other fibril types might be completely resistant. However, this is unlikely as we could not detect the disappearance of a certain type of fibril after chaperone treatment by TEM. Alternatively, the amyloid equilibrium might prevent disaggregation completion. Amyloid fibrils exist in equilibrium with monomeric species in a solution with the amyloid state highly favored. Disaggregation produces both monomeric and oligomeric species that have the ability to reaggregate over the course of the disaggregation reaction. The exact percentage of disaggregation likely depends on the kinetics of disaggregation in relation to the kinetics of (re-)aggregation and seeding by disaggregation products, which in turn is dependent on the respective fibril type. This ultimately leads to a conformation-specific disaggregation efficiency until the equilibrium is reached.

J-domain protein co-chaperones are known to confer substrate specificity to the Hsp70 machinery (Kampinga and Craig, 2010). Our data revealed that the class B J-domain proteins, DnaJB1 and DnaJB4, enabled Hsc70 to disaggregate Tau fibrils. In contrast, both of the two major cytosolic class A J-domain proteins we tested did not mediate Tau disaggregation. This is consistent with our past observations with α-synuclein fibrils (Gao et al., 2015). Class A J-domain proteins appear to be successful in prevention of aggregation of monomeric Tau (Mok et al., 2018), whereas class B J-domain proteins are capable of recognizing preformed Tau fibrils as substrates and recruiting the Hsp70 disaggregation machinery to this kind of polymeric clients. However, it remains to be addressed why DnaJB1 and DnaJB4 are so potent at supporting the disassembly of amyloid fibrils.

An intriguing difference between the disassembly reactions of α-synuclein and Tau is that Tau disaggregation did not lead to a significant increase of intermediate length fragments, but primarily produced low molecular weight species instead. Despite containing predominantly monomeric, and little oligomeric Tau species, this material had significant seeding propensity in a biosensor HEK293 cell model expressing full-length TauP301S and induced the formation of self-propagating Tau species (Fig. 6D). This implies that low molecular weight species or even monomeric Tau might be seeding-competent, which is in line with a recent study showing that fibril-derived Tau monomers exhibit seeding activity (Mirbaha et al., 2018). Hence, monomeric or small oligomeric Tau liberated from fibrils by chaperone action might still maintain a seeding-competent conformation that is different than naïve monomeric Tau.

Since amyloid structures propagate by seed-induced templated misfolding (Goedert et al., 2017), chaperone-mediated disaggregation might be detrimental *in vivo* by generating additional seeds that are able to sequester more Tau into amyloid aggregates, thereby accelerating disease progression. In support of this idea, we recently showed that the Hsp70 disaggregation machinery contributes to prion-like spreading of amyloidogenic proteins in *C. elegans* (Tittelmeier et al., 2019). Compromising the disaggregation machinery by knocking down the *C. elegans* homolog of Apg2 reduced disaggregation of α-synuclein and polyglutamine (Q_35_) aggregates, thereby decreasing their amplification and toxicity. Hence, it is tempting to speculate that the chaperone-mediated disaggregation of Tau plays a similar role in the prion-like propagation of Tau pathology throughout the brain.

Chaperone activity is commonly believed to decline during aging and in the context of neurodegenerative diseases. However, this view is too simplified. Rather, a recent study revealed that an imbalance occurs, where individual members are deregulated, with some being up- and others downregulated in the aging or diseased human brain (Brehme et al., 2014). Intriguingly, DnaJB4 gets induced in AD, PD, and HD. Since it promoted disaggregation as efficiently as DnaJB1 in our study (Fig. 4), this co-chaperone could directly link enhanced disaggregation activity to amyloid propagation in neurodegeneration.

On the other hand, Tau disaggregation mediated by chaperones is not isolated in the cellular context and might be coupled to protein degradation via the proteasome or autophagy. These pathways might be more potent in degrading smaller oligomers and monomers instead of larger fibrils, which would render disaggregation activity beneficial overall. Further studies are necessary to clarify the exact role of the Hsp70 disaggregation machinery in Tau amyloid aggregation and toxicity *in vivo*, especially because heparin-induced Tau filaments differ from those isolated from AD or Pick’s disease patients (Goedert et al., 2018; Zhang et al., 2019). Such studies will allow evaluating the therapeutic potential of the Hsp70 chaperone machinery in Tauopathies.

## Acknowledgement

We are grateful for the excellent technical assistance of Silke Druffel-Augustin, Regina Zahn, Tracy Bellande, and Audrey Coens, and for the support of the DKFZ Core Facility for Electron Microscopy (K. Richter), DKFZ Light Microscopy Core Facility (D. Krunic), and ZMBH Flow Cytometry & FACS Core Facility (M. Langlotz). We also thank Dr. Cindy Voisine and all members of the Nussbaum lab for their helpful discussion and constructive comments on the manuscript.

This study is part of the PROTEST-70 project within the EU Joint Programme - Neurodegenerative Disease Research (JPND) project. This project is supported through the following funding organizations under the aegis of JPND - www.jpnd.eu: France, Agence National de la Recherche (ANR, ANR-17-JPCD-0005-01 to R.M.); Germany, Bundesministerium für Bildung und Forschung (BMBF, 01ED1807A to B.B. and 01ED1807B to C.N-K.); The Netherlands, Netherlands Organization for Scientific Research (ZonMw - project number 733051076). Funding was also provided by Alzheimer Forschung Initiative e.V. (AFI), grant #17054 (to B.B.), the Baden-Wu□rttemberg Stiftung, BWST-ISFIII-029 (to B.B.), Centre National de la Recherche Scientifique, the Institut de France-Fondation Simone et Cino Del Duca, and the Fondation Pour La Recherche Médicale (contract DEQ. 20160334896). This work has also received support from the EU/EFPIA/Innovative Medicines Initiative 2 Joint Undertaking (IMPRiND grant No 116060). W.A.M. was supported by a Sir Henry Dale Fellowship jointly funded by the Wellcome Trust and the Royal Society (Grant Number 206248/Z/17/Z). E.N. was supported by PhD fellowships from the Helmholtz International Graduate School for Cancer Research (DKFZ), the Chica and Heinz Schaller Foundation, and the Studienstiftung des deutschen Volkes.

## Contributions

Conceptualization, E.N., T.R.J., B.B., and C.N.-K.;

Methodology, E.N., A.W., K.M., L.B., T.K., W.A.M., R.M., A.M., and C.N.-K.;

Investigation, E.N.;

Formal Analysis, E.N.;

Resources, E.N., A.W., K.M., L.B., T.K., W.A.M., and R.M.;

Writing – Original Draft, E.N. and C.N.-K.;

Writing – Review and Editing, E.N., A.W., T.K., H.H.K., W.A.M., T.R.J., R.M., A.M., B.B., and C.N.-K.;

Supervision, W.A.M., T.R.J., R.M., B.B., and C.N.-K.;

Visualization, E.N. and C.N.-K.;

Funding Acquisition, A.W., H.H.K., W.A.M., T.R.J., R.M., B.B., and C.N.-K.;

## Material and Methods

All chemicals were purchased from Sigma-Aldrich or Carl Roth unless stated otherwise.

### Purification of recombinant proteins

The six isoforms of full-length human Tau, e.g. 0N3R, 0N4R, 1N3R, 1N4R, 2N3R and 2N4R Tau, were expressed and purified as described previously (Tardivel et al., 2016). Tau protein concentration was determined spectrophotometrically using an extinction coefficient at 280 nm of 7450 M^−1^cm^−1^. Aliquots of pure Tau isoforms (100 *µ*M) were stored at −80 °C.

Monomeric Tau was used as a control in several experiments of this study. Aliquots were stored at −80 °C. In order to remove aggregates that might have formed during the freeze-thaw process, samples were centrifuged at least at 100 000 g immediately before using the supernatant for any experiment.

The human chaperones Hsc70, DnaJB1, DnaJA1, DnaJA2, and Apg2 were purified as previously published (Andreasson et al., 2008; Gao et al., 2015; Rampelt et al., 2012). Briefly, N-terminally His6-Sumo tagged proteins were expressed in BL21 *E. coli* (DE3) and affinity purified using Protino Ni-NTA Agarose (Macherey-Nagel). Subsequently, the tag was cleaved off by Ulp-1 digest and both the His6-Sumo-tag and the His tagged Ulp1 were removed by a second Ni^2+^ affinity purification step. The proteins were further purified by size exclusion chromatography on a Superdex200 16/60 column (GE Healthcare).

The human DnaJB4 DNA sequence (The ORFeome Collaboration (2016), DKFZ) was cloned into a pCool6 vector with an N-terminal His6-Sumo tag generated previously (Ho et al., 2019) and expressed at 16 °C. Further purification steps were performed following the protocol stated above. Aliquoted proteins were stored at −80 °C.

### *In vitro* aggregation of recombinant Tau

Fibrillation of the six Tau isoforms was achieved at 40 μM in the presence of 10 μM heparin by shaking 0.5 ml solution aliquots at 37 °C in an Eppendorf Thermomixer set at 600 rpm for 4 days. At steady state, an aliquot from each assembling reaction was spun for 35 min at 20 °C and 50 000 rpm (113 000 g) and the amount of Tau in the supernatant assessed spectrophotometrically and by SDS-PAGE to further demonstrate assembly completion. The amount of fibrillar Tau was estimated by subtraction of the soluble fraction remaining after centrifugation from the initial concentration.

### *In vitro* disaggregation assay

*In vitro* disaggregation reactions were performed as previously described (Gao et al., 2015) with minor adjustments. Briefly, recombinant Tau fibrils aggregated *in vitro* (2 *µ*M) or Sarkosyl-insoluble material extracted from seeded TauP301S-Venus HEK293 cells were incubated with the disaggregation machinery (Hsc70 (4 *µ*M), DnaJB1 (2 *µ*M), and Apg2 (0.2 *µ*M)) in disaggregation buffer (50 mM Hepes-KOH (pH 7.5), 50 mM KCl, 5 mM MgCl_2_, and 2 mM DTT) at 30 °C for the indicated timespans. For “+ATP” conditions 2 mM ATP and an ATP regeneration system (4.5 mM phosphoenolpyruvate, 20 ng/ml pyruvate kinase (Sigma-Aldrich)) were added to the reaction. In “-ATP” conditions both ATP and the ATP regeneration system were omitted. After the indicated incubation times, samples were centrifuged for 30 min at 20 000 g, or 337 000 g, 4 °C. Tau levels in supernatant (S) or pellet (P) fractions were subsequently analyzed by SDS-PAGE and immunoblotting.

### ThT disaggregation assay

For kinetic analysis of the disaggregation reaction, 50 *µ*l samples were prepared as described above, but including 20 *µ*M Thioflavin T (ThT). Buffer without Tau fibrils or chaperones was used as blank control. The measurement was performed in sealed (Microseal B Adhesive Sealer, BioRad) black 96-well clear bottom plates (flat bottom, non-binding surface, Corning) at 30 °C using a FLUOstar Omega plate reader (BMG Labtech). ThT fluorescence was recorded by bottom reading every 5 min with excitation/emission wavelengths set to 440/480 nm. Before each acquisition cycle the plate was shaken for 10 s at 300 rpm. Data was normalized to the untreated fibril control and chaperone -ATP condition for each time point.

### Rate-zonal centrifugation

Continuous 5-45% sucrose gradients (in 50 mM Hepes-KOH (pH 7.5), 50 mM KCl, 5 mM MgCl_2_) were formed in 12 ml open-top polyclear centrifuge tubes (Seton Scientific) on the Gradient Station *ip* (BioComp Instruments). Tau fibrils were incubated with the disaggregation machinery at 30 °C and after 0.5 h and 4 h aliquots were taken from the reaction mix and applied to the sucrose gradients. Centrifugation was performed at 22 °C for 3 h at 217 874 g using a SW 40 Ti rotor (Beckman Coulter). Fractions of 600 *µ*l were carefully collected manually. During the manual fractionation a thin film of low-density sucrose always remained on top of the gradients which was then collected with the pellet fraction. Therefore, the protein amount in the last fraction is slightly overestimated. The pellet was resuspended in equal volumes 1x Laemmli in PBS. Samples were analyzed by SDS-PAGE and immunoblotting. The amount of Tau in each fraction as percent of total Tau across the whole gradient was quantified using the Image Studio Lite software (LI-COR Biosciences).

### SDS-PAGE and immunoblotting

Samples were run on either 4-20% Express Plus PAGEs in Tris-MOPS-SDS running buffer (GenScript), 10% Criterion TGX gels (BioRad) in Tris-glycine-SDS running buffer or 3-8% Criterion XT Tris-acetate gels (BioRad) in Tris-acetate-SDS running buffer. Proteins were transferred to PVDF membranes (Trans-Blot Turbo RTA Transfer Kit, BioRad) using the Trans-Blot Turbo Transfer System (BioRad) and immunoblotted with the Tau antibody A-10 (1:1 000 - 10 000, mouse, sc-390476, Santa Cruz Biotechnology) or anti-GFP (1:10 000, mouse, MMS-118P, Covance). An alkaline phosphatase-coupled secondary antibody (Vector Laboratories) together with ECF substrate (GE Healthcare Life Sciences) was used for development. The blots were imaged on an ImageQuant LAS-4000 (FUJIFILM Co.). Densitometric quantification of the signals was performed with the Image Studio Lite software (LI-COR Biosciences).

### Negative stain electron microscopy

Tau fibrils alone or treated with chaperones in the absence or presence of ATP and ATP regenerating system were diluted in PBS or disaggregation buffer, respectively and pipetted onto carbon-coated copper grids (Plano GmbH). Samples were allowed to absorb for 1 min before washing twice with 10 *µ*l water for 1 min. Negative stain was achieved by incubation with 2% (w/v) aqueous uranyl acetate for 1 min. Excess solution was removed by blotting the grids carefully on filter paper before imaging on an EM-900 or an EM-910 electron microscope (Zeiss) with an accelerating voltage of 80 kV.

### Cell culture

The HEK293 cell line expressing 0N4R TauP301S-Venus generated by McEwan et al. (2017) was cultured in Dulbecco’s Modified Eagle Medium (DMEM), high glucose, GlutaMAX Supplement, pyruvate (Gibco) supplemented with 10% fetal calf serum (FCS) (Gibco), 100 IU/ml penicillin, and 100 mg/ml streptomycin (Gibco) at 37 °C and 5% CO_2_. Cells were regularly tested for mycoplasma contamination.

### Generation of enriched seeded pool of TauP301S-Venus HEK293 cells

The naïve TauP301S-Venus HEK293 cell line was treated with preformed 1N4R Tau fibrils as published (McEwan et al., 2017). Briefly, cells were plated in a 6-well plate in Opti-MEM Reduced Serum Medium, GlutaMAX Supplement (Gibco). The next day the cells were treated with 100 nM preformed Tau fibrils and 10 *µ*l Lipofectamine2000 (Invitrogen) diluted in Opti-MEM. After 1 h incubation time, equal volumes of complete DMEM were added. Three days later, the cells were washed and resuspended in cell sorting buffer (1x PBS + 0.8 mM EDTA, 0.5% (v/v) FCS) before carefully passing the cells through a 35 *µ*m cell strainer (Corning). 7500 foci-containing cells were sorted based on the intracellular distribution of Venus fluorescence (concentrated intensity signal vs. diffuse distribution) on FACS Aria IIIu (Becton Dickinson) with a 530/30 nm filter at the ZMBH FACS facility. The derived cell population was collected in complete DMEM and expanded until stocks could be frozen. The presence of TauP301S-Venus foci before and after freezing/thawing was confirmed by fluorescence microscopy.

### Extraction of Sarkosyl-insoluble material from seeded TauP301S-Venus HEK293 cells

HEK293 cells propagating aggregated TauP301S-Venus were harvested by snap freezing. The cell pellet was resuspended in cold extraction buffer (10 mM Tris pH 7.5, 2 mM NaV, 50 mM NaF, 50 mM β-glycerophosphate, PhosSTOP Phosphatase inhibitor (Roche), cOmplete EDTA-free Protease Inhibitor Cocktail (Roche), 100 mM NaCl) and sonicated shortly. Cell debris was removed by centrifugation at 1 000 g, 4 °C for 1 min. The cleared lysate was centrifuged at 337 000 g, 4°C, for 15 min. The resulting pellet was resuspended in 100 *µ*l extraction buffer with 1% (w/v) Sarkosyl, sonicated again and then incubated for 1 h at 22 °C, 700 rpm, to extract the Sarkosyl-soluble fraction before repeating the centrifugation step. The Sarkosyl-insoluble pellet was resuspended in disaggregation buffer with protease inhibitors (50 mM Hepes-KOH (pH 7.5), 50 mM KCl, 5 mM MgCl_2_, 2 mM DTT, 2 mM NaV, 50 mM NaF, 50 mM β-glycerophosphate, PhosSTOP Phosphatase inhibitor (Roche), cOmplete EDTA-free Protease Inhibitor Cocktail (Roche)), shortly sonicated and centrifuged to remove the remaining detergent. Finally, the pellet was resuspended in disaggregation buffer with protease inhibitors and stored at 4 °C.

### *In vivo* seeding assay

To test the seeding capacity of Tau liberated by the disaggregation machinery, disaggregation reactions were performed as described above. In order to obtain the fraction of Tau that was liberated by chaperone action, differential centrifugation steps first at 20 000 g and then at 337 000 g were performed, both for 30 min, 4 °C. After ultracentrifugation only the upper two thirds of the supernatant were carefully collected thus avoiding disturbing the pelleted material. Samples of all fractions were subjected to SDS-PAGE and immunoblotting to confirm successful differential centrifugation.

The seeding assay with the biosensor TauP301S-Venus HEK293 was performed following the protocol for liposome-mediated transduction by McEwan et al. (2017). Briefly, 50 000 cells were seeded per 24-well on Poly-L lysine-coated coverslips in 300 *µ*l Opti-MEM Reduced Serum Medium, GlutaMAX Supplement (Gibco). The next day, the cells were treated with 25 *µ*l of the 337 000 g supernatants of the disaggregation reactions. The samples were mixed with 2.5 *µ*l Lipofectamine2000 (Invitrogen) in 200 *µ*l OptiMEM and added to the cells. After 1 h treatment, the seeding reaction was stopped by adding 500 *µ*l complete DMEM to each well. 24 h later, the cells were fixed in 4% PFA in PBS for 30 min. After washing, the cells were incubated with 0.1 *µ*g/ml DAPI in PBS and mounted in Vectashield (Vector Laboratories) for fluorescence microscopy.

In order to monitor the propagation of TauP301S-Venus foci over time, the cells were passaged for 27 days (6 passages) after seeding and imaged regularly with a Leica DM IL LED system equipped with a HI PLAN I Phase 2 40x/0.50 (Leica) objective lens.

### Microscopy and image analysis of fixed cells

Fixed TauP301S-Venus HEK293 cells were imaged using a Zeiss Cell Observer equipped with a Plan-Apochromat 20x/0.8 M27 (Zeiss) objective lens. For each image three z-stacks at intervals of 20 *µ*m were acquired in order to capture all foci within a cell. Semi-automated image analysis was performed using Fiji (Schindelin et al., 2012) and a macro for counting nuclei and for foci identification was developed together with the DKFZ Light Microscopy Core Facility. Afterwards, the foci were manually assigned to single cells in order to calculate the percentage of cells in which Tau aggregation was seeded. Per replicate and condition at least 288 cells were analyzed.

### Statistical analysis

The statistical analysis was performed using GraphPad Prism (GraphPad Software, Version 6). For each dataset the sample size (n), p-values, and the statistical test, which was applied, are indicated in the corresponding figure legend.

All data are shown as mean values ± standard error of the mean (SEM) with the following significance levels: non-significant (ns), p > 0.05, ^*^p ≤ 0.05, ^**^p ≤ 0.01, and ^***^p ≤ 0.001.

