## Supplemental Information for "Disassembly of Tau fibrils by the human Hsp70 disaggregation machinery generates small seeding-competent species"

#### Supplementary Figure Legends

**Figure S1. Alternative splicing of the human MAPT gene gives rise to six different Tau isoforms.**

The different isoforms are named based on their domain structures (left). Alternative splicing events in the N-terminal region (N1 and N2, red), give rise to the three different variants 2N, 1N, and 0N. The microtubule-binding domain (MTBD) consists of either three (3R) or four (4R) repeat domains (blue) and forms the major part of the fibril core in Tau amyloid assemblies. Alternative splicing of exon 10 (R2, light blue) within the MTBD determines the R variant (3R or 4R). The C-terminal region (C) is identical in all six human Tau variants.

**Figure S2. Class B J-domain proteins mediate Tau disaggregation.**

**(A)** Fibrils of all six Tau isoforms were treated with Hsc70, Apg2, and DnaJA2 in the presence or absence of ATP. S and P fractions were separated by centrifugation and Tau levels were analyzed by immunoblotting. DnaJA2 did not mediate Tau disaggregation *in vitro.*

**(B)** Sequence alignment of indicated J-domain proteins. Class B J-domain proteins DnaJB1 and DnaJB4 share a sequence identity of 65.7%.

**(C)** Phylogram of the class A and class B J-domain proteins analyzed in this study.

**Figure S3. Characterization of monomeric and fibrillar Tau without the addition of chaperones.**

Monomeric Tau or fibrils were centrifuged over a 5-45% sucrose gradient. Fractions were collected manually, and the Tau content was analyzed by immunoblotting. Monomeric Tau was detected in the first two fractions, while Tau fibrils migrated to high-density fractions. Fractionating a gradient loaded with monomeric 1N4R Tau without previous centrifugation (no centrifugation, NC) demonstrated that minor amounts of monomeric Tau could be detected in the pellet fraction (marked with *) for technical reasons.

**Figure S4. No accumulation of smaller Tau fragments after disaggregation.**

Tau fibrils were treated for 20 h with the human Hsp70 disaggregation machinery in the presence or absence of ATP. Negative stained samples were analyzed by TEM. Scale bar = 500 nm.

**Figure S5. Tau disaggregation generates monomeric and small oligomeric seeding-competent species.**

**(A)** Maximum intensity projections of fluorescence microscopy z-stacks of TauP301S-Venus HEK293 cells seeded with the 337 000 g supernatant fraction of the buffer control after the disaggregation reaction or naïve monomeric Tau. Scale bar = 50 µm. Neither caused foci formation in the TauP301S-Venus HEK293 cell culture model.

**(B)** Epi fluorescence microscopy of TauP301S-Venus HEK293 cells treated with the 337 000 g supernatant fractions after the disaggregation reaction. The cells were imaged 2 days and 27 days after the treatment. Scale bar = 50 µm.

**Supplementary Figure S1**

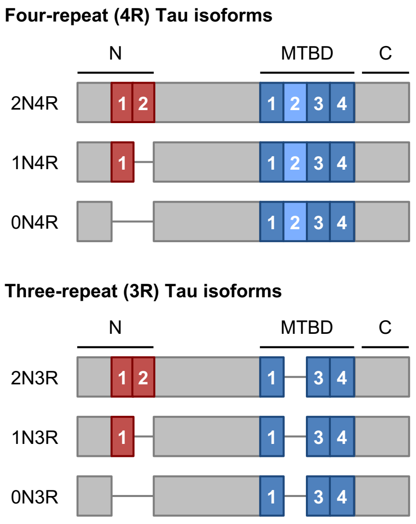

### Supplementary Figure S2

A

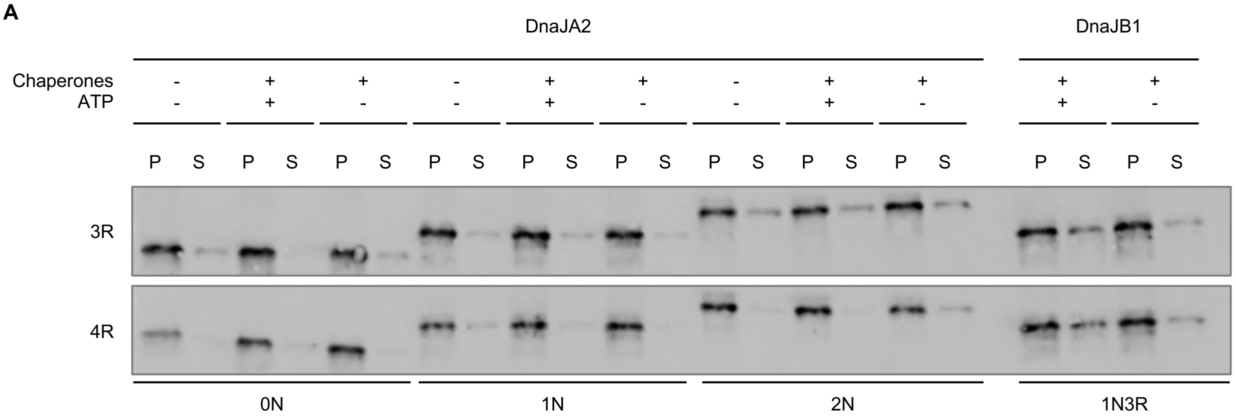


B

```
DNJB1|P25685      ----MGKDYYQTLGLARGASDEEIKRAYRRQALRYHPDKNKEPGAEEKFKEIAEAYDVLS  56
DNJB4|Q9UDY4      ----MGKDYYCILGIEKGASDEDIKKAYRKQALKFHPDKNKSPQAEEKFKEVAEAYEVLS  56
DNJA1|P31689      --MVKETTYYDVLGVKPNATQEELKKAYRKLALKYHPDKN--PNEGEKFKQISQAYEVLS  56
DNJA2|O60884      MANVADTKLYDILGVPPGASENELKKAYRKLAKEYHPDKN--PNAGDKFKEISFAYEVLS  58
                        .  *  **:  .*:::::*:***: * .:*****  *   :***::: **:***

DNJB1|P25685      DPRKREIFDRYGEEGLKGSGPSGGSGGGANGTSFSYTFHGDPHAMFAEFFGGRNPFDTFF  116
DNJB4|Q9UDY4      DPKKREIYDQFGEEGLKGGA----GGTDGQGGTFRYTFHGDPHATFAAFFGGSNPFEIFF  112
DNJA1|P31689      DAKKRELYDKGGEQAIKEGGAGGGF--------------GSPMDIFDMFFGGGG------  96
DNJA2|O60884      NPEKRELYDRYGEQGLREGSGGGG----------------GMDDIFSHIFGGGL------  96
                  : .***::*: **:.:: ..                    .    *  :***

DNJB1|P25685      GQRNGEEGMDIDDPFSGFPMGMGGFTNVNFGRSRSAQEPARKKQDPPVTHDLRVSLEEIY  176
DNJB4|Q9UDY4      GRRMGGGRDSEEMEIDGDPFSAFGFSMNGYPRDRNSVGPSRLKQDPPVIHELRVSLEEIY  172
DNJA1|P31689      -------------------------------------RMQRERRGKNVVHQLSVTLEDLY  119
DNJA2|O60884      ----------------------FGFMGNQ------SRSRNGRRRGEDMMHPLKVSLEDLY  128
                                                            ::.  : * * *:**::*

DNJB1|P25685      SGCTKKMKISHKRL----------------------------------------------  190
DNJB4|Q9UDY4      SGCTKRMKISRKRL----------------------------------------------  186
DNJA1|P31689      NGATRKLALQKNVICDKCEGRGGKKGAVECCPNCRGTGMQIRIHQIGPGMVQQIQSVCME  179
DNJA2|O60884      NGKTTKLQLSKNVLCSACSGQGGKSGAVQKCSACRGRGVRIMIRQLAPGMVQQMQSVCSD  188
                  .* * :: :.:: :

DNJB1|P25685      ---------------NPDGKSIRNEDKILTIEVKKGWKEGTKITFPKEGDQTSNNIPADI  235
DNJB4|Q9UDY4      ---------------NADGRSYRSEDKILTIEIKKGWKEGTKITFPREGDETPNSIPADI  231
DNJA1|P31689      CQGHGERISPKDRCKSCNGRKIVREKKILEVHIDKGMKDGQKITFHGEGDQEPGLEPGDI  239
DNJA2|O60884      CNGEGEVINEKDRCKKCEGKKVIKEVKILEVHVDKGMKHGQRITFTGEADQAPGVEPGDI  248
                                 . :*:.   * *** :.:.** *.* :***  *.*:  .  *.**

DNJB1|P25685      VFVLKDKPHNIFKRDGSDVIYPARISLREALCGCTVNVPTLDGRTIPVVF--KDVIRPGM  293
DNJB4|Q9UDY4      VFIIKDKDHPKFKRDGSNIIYTAKISLREALCGCSINVPTLDGRNIPMSV--NDIVKPGM  289
DNJA1|P31689      IIVLDQKDHAVFTRRGEDLFMCMDIQLVEALCGFQKPISTLDNRTIVITSHPGQIVKHGD  299
DNJA2|O60884      VLLLQEKEHEVFQRDGNDLHMTYKIGLVEALCGFQFTFKHLDGRQIVVKYPPGKVIEPGC  308
                  ::::.:* *  * * *.::     * * *****    .  **.* * :     .::. *

DNJB1|P25685      RRKVPGEGLPLPKTPEKRGDLIIEFEVIFPERIP--QTSRTVLEQVLPI-----------  340
DNJB4|Q9UDY4      RRRIIGYGLPFPKNPDQRGDLLIEFEVSFPDTIS--SSSKEVLRKHLPAS----------  337
DNJA1|P31689      IKCVLNEGMPIYRRPYEKGRLIIEFKVNFPENGFLSPDKLSLLEKLLPERKEVEET-DEM  358
DNJA2|O60884      VRVVRGEGMPQYRNPFEKGDLYIKFDVQFPENNWINPDKLSELEDLLPSRPEVPNIIGET  368
                   : : . *:*  : * ::* * *:*.* **:       .   *.. **

DNJB1|P25685      ------------------------------------------  340
DNJB4|Q9UDY4      ------------------------------------------  337
DNJA1|P31689      DQVELVDFDPNQ-----ERRRHYNGEAYEDDEHHPRGGVQCQTS  397
DNJA2|O60884      EEVELQEFDSTRGSGGGQRREAYNDSSDEESSSHHGPGVQCAHQ  412
```

C

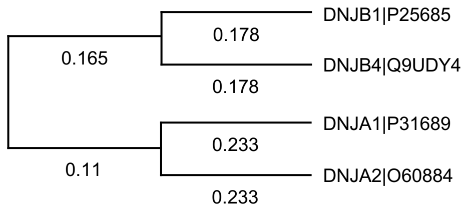

#### Supplementary Figure S3

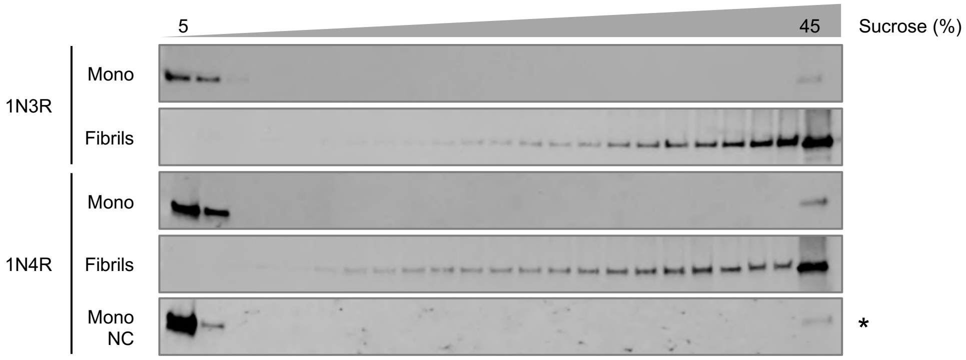

### Supplementary Figure S4

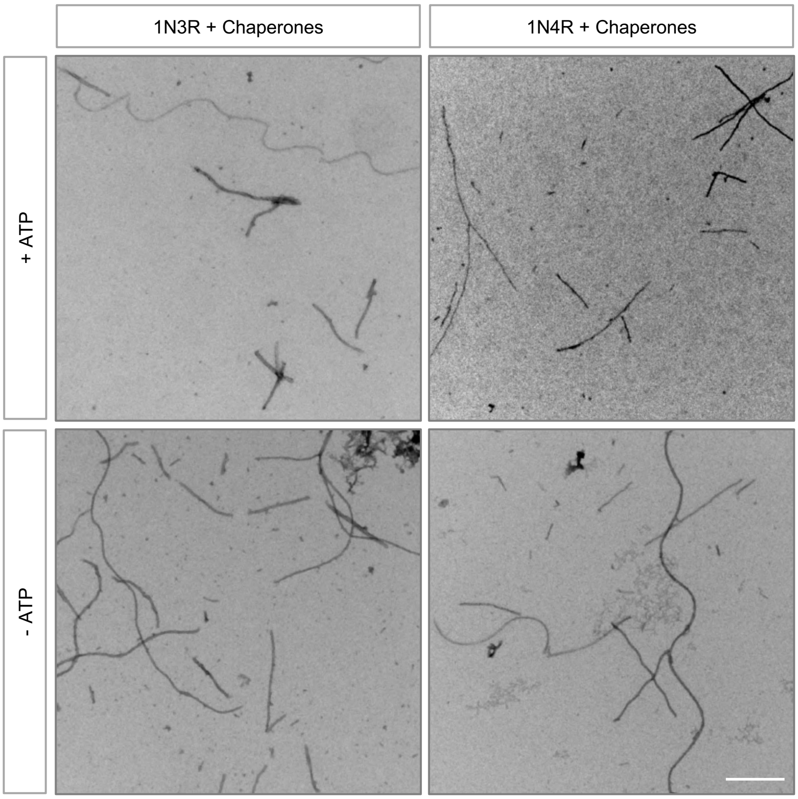

#### Supplementary Figure S5

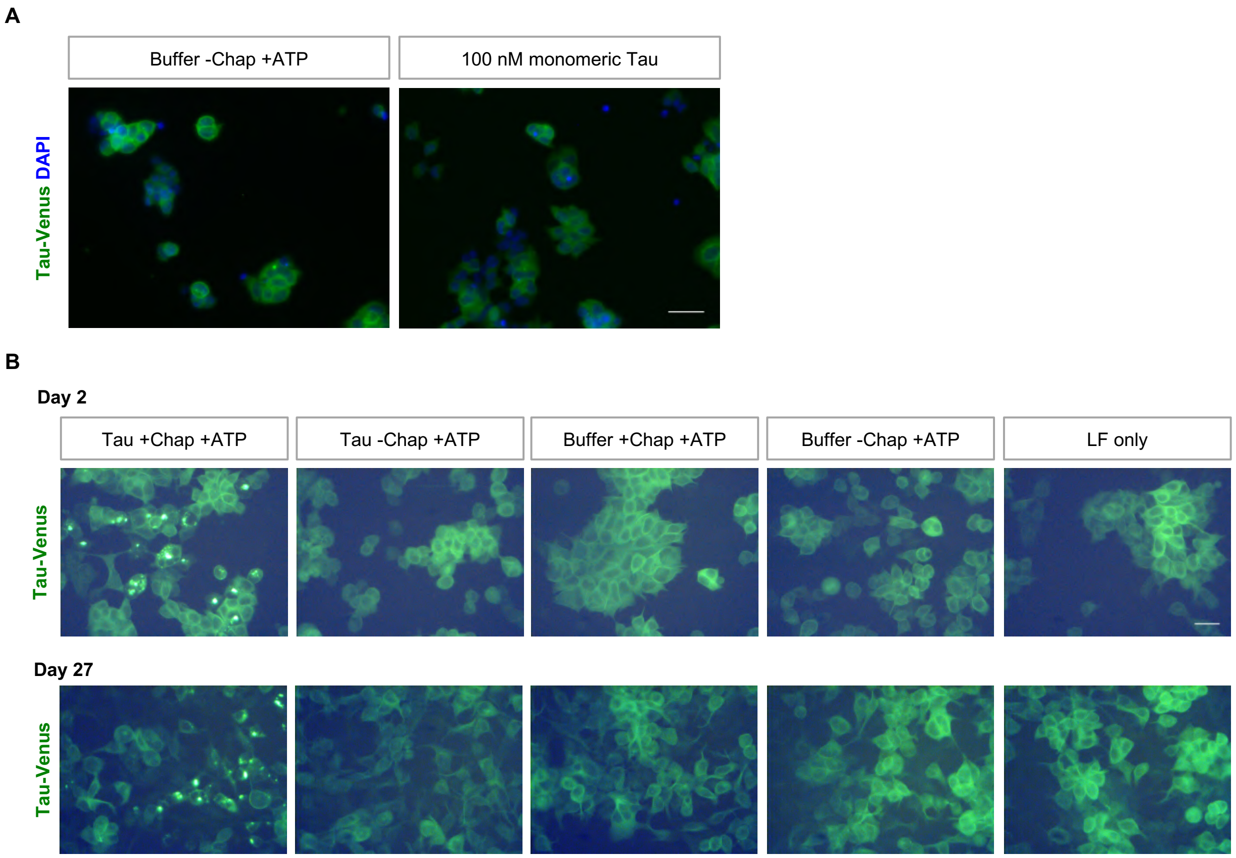
